# Mapping the loss of flagellar motility across the tree of life

**DOI:** 10.1101/2024.12.05.626484

**Authors:** Jamiema Sara Philip, Sehhaj Grewal, Jacob Scadden, Caroline Puente-Lelievre, Nicholas J. Matzke, Luke McNally, Matthew AB Baker

## Abstract

Bacterial swimming is mostly powered by the bacterial flagellar motor and the number of proteins involved in the flagellar motor can vary. Quantifying the proteins present in flagellar motors from a range of species delivers insight into how motility has changed throughout history and provides a platform for estimating from its genome whether a species is likely to be motile. We conducted sequence and structural homology searches for 54 flagellar pathway proteins across 11,365 bacterial genomes and developed a classifier with up to 95% accuracy that could predict whether a strain was motile or not. We then mapped the evolution of flagellar motility across the GTDB bacterial tree of life. We confirmed that the last common bacterial ancestor had flagellar motility and that the rate of loss of this motility was four-fold higher than the rate of gain. We showed that the presence of filament protein homologues was highly phylogenetically correlated with motility and that all species classified as motile contained at least one filament homologue. We calculated the rate of gain and loss for each flagellar protein and that the filament protein FliC was highly correlated with motility across the tree of life. We then measured the correlation of each flagellar motor protein with FliC and showed that the filament, rotor, and rod and hook proteins were all highly correlated with FliC, and thus with motility. We calculated the differential rates of gain and loss for each flagellar protein and quantified which genomes encoded for partial sets of flagellar proteins, indicating potential pathways by which motility could be lost. Overall, we show that filament, rod and hook and rotor proteins are conserved when flagellar motility is preserved and that the presence or absence of a FliC homologue is a good, simple predictor of whether or not a species has flagellar motility.

## Introduction

Flagellar motility is widespread among bacteria, with the flagellar apparatus exhibiting diverse structures and arrangements across species [1]. Despite these structural differences, all flagellated bacteria share a common rotary nanomachine known as the bacterial flagellar motor (BFM), which generates thrust through coupling ion flux to torque generation [2]. The motor has conserved core structures with divergent structures surrounding them [3–5]. These additional components and modifications in the motor structure reveal its adaptation to diverse environments, increasing bacterial fitness [6–8].

Flagellar assembly and function are regulated by a complex network of genes that have evolved considerably across different bacterial lineages [8]. Although much is known about the structure and function of the BFM, little is known about the evolutionary origins of the gene clusters that define these diverse organelles. Early work identified 20 essential genes for flagellar assembly and function in *Escherichia coli* [9]. Over time, research revealed the involvement of nearly 40 genes in *E. coli* and *Salmonella enterica* [10], with some species like *Vibrio parahaemolyticus* possessing dual flagellar systems—each expressed as distinct gene sets comprising around 50 polar and 40 lateral flagellar genes [11]. A broader comparative study of 41 flagellated bacterial species identified a core set of 24 structural genes thought to have been present in the last common bacterial ancestor (LCBA) [8]. Subsequent rooting of the bacterial tree and reconstruction of the eubacterial ancestor further verified that the LCBA was flagellated [12].

Although flagellar motors and motility mechanisms have been extensively studied in model species, our understanding of motility across diverse bacterial groups remains limited. Recent research in Firmicutes reveals that flagellar motility and cell shape evolved independently challenging previous assumptions [13]. Moreover, flagellar motility which was an ancestral trait in *Chlamydiae* [14] and *Dehalococcoidia* [15], has been lost in various lineages, reflecting its adaptation to various environmental pressures. For instance, in *Rhodanobacter*, flagellar loss is associated with stress adaptation, such as enhanced biofilm formation [16]. These findings highlight the need to expand our perspective on flagellar evolution beyond model species and recognise the complexity and diversity of motility across bacterial phyla. Efforts have been made to quantify flagellar genes present in soil communities, but these were limited to specific species from a single site [17] making it difficult to generalise findings to other bacterial communities. A genome-based model for predicting flagellar motility was developed for soil bacterial communities [18], specifically *Actinobacteriota, Firmicutes*, and *Proteobacteria*, but this has not been extended to other phyla.

Sequence-based approaches struggle to capture flagellar protein-encoding genes across diverse species due to low sequence homology [8]. Protein structure searches offer greater sensitivity in detecting homology and resolving deep evolutionary relationships [19]. Although structural data exists for many flagellar components, including the basal body, filament, export apparatus, and regulatory proteins, some gaps remain particularly in understudied species with additional proteins [20].

Here, we utilized a dataset of 11,365 complete and non-redundant bacterial genomes from the PATRIC database, ensuring representation across diverse bacterial taxa. We employed a combination of sequence and structure-based methods to identify and quantify flagellar proteins within these genomes. Our analysis revealed a bimodal distribution in the number of different flagellar proteins that were present across the species. This allowed us to develop a classifier that predicted whether a genome was motile or non-motile based on the overall number of flagellar proteins that were present. Our classifier predictions were further validated through comparsion with phenotypic data from published reports [13, 21]. By integrating structural homology with sequence-based searches we reduced the number of false positives and improved the detection of distant flagellar homologues. Lastly, we reconstructed motility as a trait across a microbial species tree to determine where motility was lost across this tree, and to measure the individual rates of protein gain and loss, as well as the correlation of protein loss with the loss of motility.

## Materials and Methods

### Data collection

We downloaded all proteomes from 11,365 complete and non-redundant bacterial genomes from the PATRIC database (downloaded August 2023)[22](Supplementary Table S1). As a core set of flagellar proteins for subsequent querying against the database, 54 proteins from the flagellar assembly reference pathway [*map03040*] of the Kyoto Encyclopaedia or Genes and Genomes (KEGG) database[23] were selected. We assembled query datasets of flagellar proteins from six model species: *Escherichia coli, Bacillus subtilis, Pseudomonas aeruginosa, Vibrio alginolyticus, Shewanella oneidensis*, and *Vibrio alginolyticus*. Flagellar proteins such as motC, motD, and flgQ, which are found absent in the above species but present in other model organisms such as *Sinorhizobium meliloti* and *Campylobacter jejuni*, were retrieved from KEGG and added to query datasets.

### Homologue identification

A total of 308 representative sequences of 54 key flagellar proteins assembled from the above species were used to identify homologs in complete genomes (Supplementary Table S2). Homologues were identified using default settings [*jackhmmer --noali -A <alignment_output> --tblout <table_output> -o <output_file> <query_file> <database_file>]* for iterative searches with jackhmmer with criteria evalue 1E–10 [24]. To further remove false positives, we utilised additional steps, including length filtering and structural information (Fig. 1A). We defined the length range for each flagellar protein based on the min and max of representative query sequences and excluded sequences longer than this range. Approximately 2% of sequences were removed due to length (Supplementary Table S3).

**Fig. 1:**
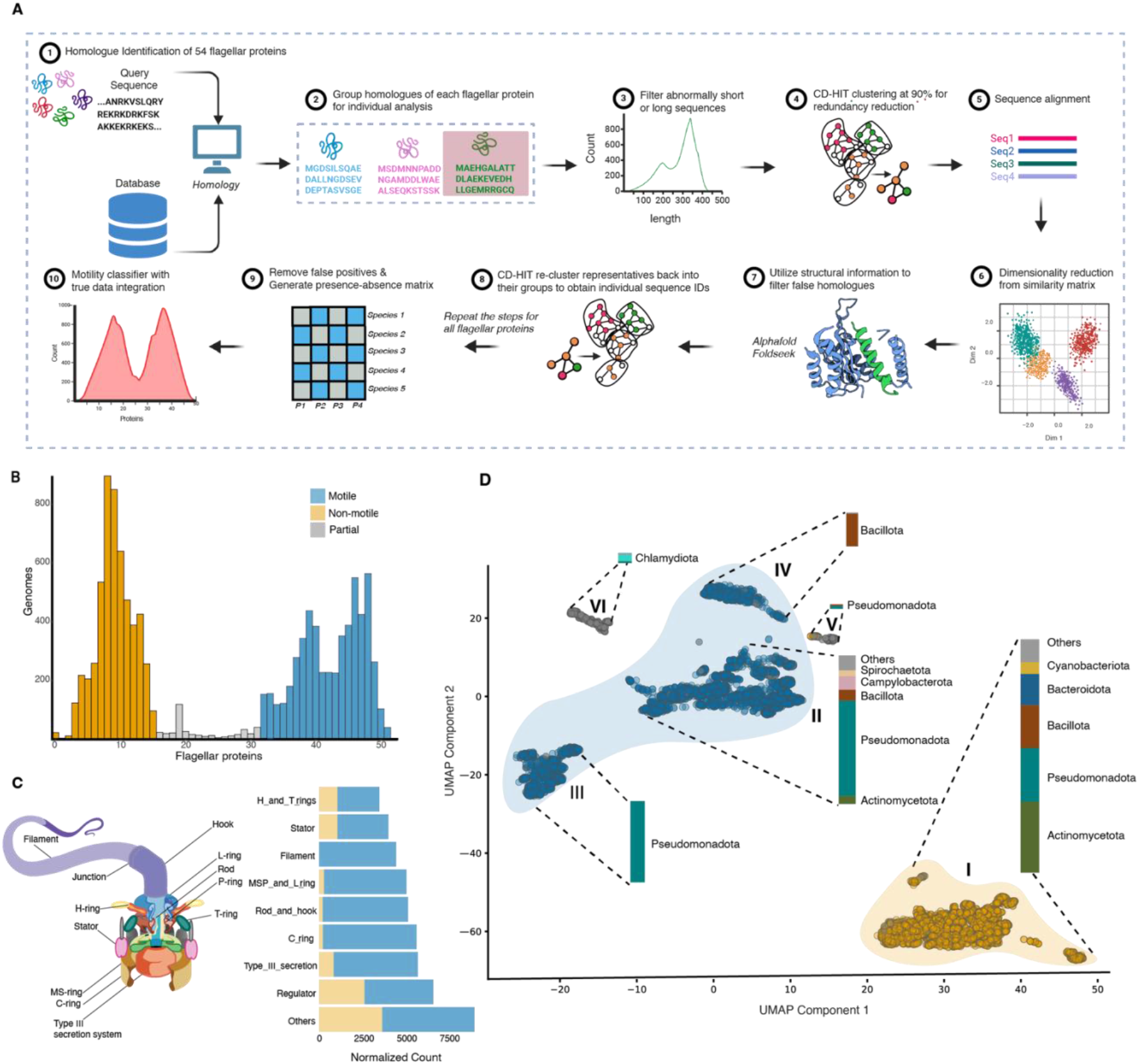
Bimodal distribution of flagellar protein number across genomes. (A) Schematic overview of the flagellar protein homologue search in bacterial genomes. The query proteins assembled from six bacterial species were searched against the PATRIC database. The sequence homologues were further processed for downstream analysis such as length filtering and structural similarity checks to improve the confidence of the homology search. (B) The density plot depicts the bimodal distribution of flagellar proteins across 11,365 genomes. Genomes with ≤ 15 flagellar proteins were categorised as non-motile, whereas those with ≥ 32 were considered motile. (C) Schematic of the BFM indicating different substructures of the motor (left) and horizontal stacked bar plot (right) showing the normalised counts of flagellar protein homologues in motile and non-motile groups (orange: non-motile; blue: motile). Counts are normalised across each class or substructure by dividing by the number of query proteins in each class. Others corresponds to the transporters and sigma factors involved in flagellar pathways. (D) UMAP visualisation of 11,365 genomes clustered based on the presence-absence matrix of flagellar proteins with DBSCAN clustering. Each dot represents a genome, with colors indicating motility classification from Figure 1B: blue for motile (genomes with ≥ 32 flagellar proteins), orange for non-motile (genomes with ≤15 flagellar proteins), and grey for genomes that have between 15 and 32 flagellar proteins, which are classified as partial protein sets. Stacked bar plots placed on each cluster depict the relative abundance of major bacterial phyla within those clusters.

Homologous sequences for each protein were clustered at 90% identity using CD-HIT [25], and representative sequences were aligned using MAFFT [26] with default parameters. These alignments were then used to generate a distance matrix and perform PCA for dimensionality reduction. We selected the ten most distant points from the centre of the PCA plot, calculated by the mean, for further structural comparison to our query proteins. Our approach was to test whether the ten most distant protein sequences for structural homology as a measure of whether the overall set of sequences was representative of homologues with similar function, involved in flagellar motility. We first predicted structures for these 10 with AlphaFold2 [27] and then used Foldseek [28] to compare each of these 10 predicted structures with the original target query structures. Predicted structures were defined as structural homologues if the Foldseek Probability was equal to one. Otherwise, these were rejected as homologues for the target proteins.

Proteins FliG and FliI had no structural homologues for the 10 most distant sequences in the PCA sequence space. For these two cases, we tested a further 10 and again found no structural homologues. To better separate clusters for FliG and FliI we then employed UMAP (Uniform Manifold Approximation and Projection) for dimensionality reduction along with DBSCAN (Density-Based Spatial Clustering of Applications with Noise) [29]. UMAP is a nonlinear dimensionality reduction method that preserves both local and global structures [30]. We optimised UMAP parameters (n_neighbors=200 and min_dist=0.3), based on clear separation between and preservation of global structures (Supplementary Fig. S1). We searched within clusters for available AlphaFold structures and used Foldseek to determine whether they represented true FliG/FliI clusters based on Foldseek probability. Clusters were excluded as query hits if the cluster contained no structures that were structural homologs of a query protein. When structures were excluded, all sequences that shared the original sequence cluster from CD-HIT were also excluded.

### Motility classifier

For determination of whether a specific species was motile or not we listed each bacterial species and the total number of flagellar query proteins for which a hit had been found. Histogram plots were generated using the ggplot2 package to examine distribution patterns. The bimodal distribution revealed two distinct peaks, which were classified into motile and non-motile clusters. The number of flagellar proteins (N_FP_, x-axis, Fig. 1B) at which these clusters separate was determined visually from the distribution and species were declared non motile species for N_FP_ ≤ 15, motile for N_FP_ ≥ 32, with species declared as partial protein sets for 15 < N_FP_ < 32. UMAP visualization was employed to analyze the presence-absence matrix of flagellar proteins across the genomes to identify any clear clusters separating motile from non-motile genomes (Fig. 1D).

[additional methods paragraph here stating exactly how the normalisation/abundance was done for Fig. 2]

**Fig. 2:**
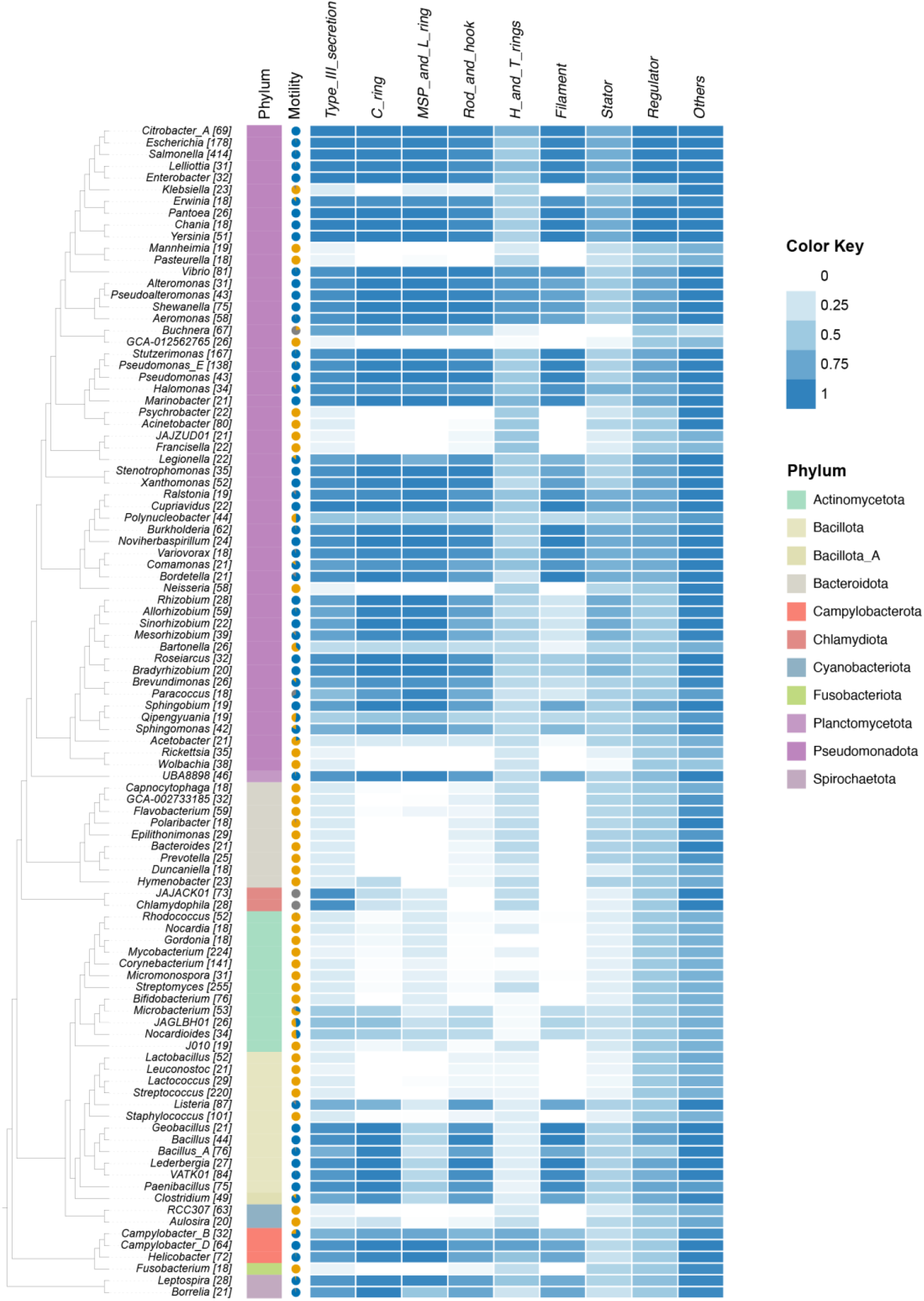
Flagellar protein distribution across the reference GTDB bacterial tree. The heatmap shows the normalised abundance of flagellar protein homologues across the most prevalent genera in the dataset (18 < N < 414). Presence-absence counts were aggregated at the genus level and normalised based on both the number of strains within each genus and the number of proteins in each functional class (Fig. 1C). The adjacent cladogram illustrates the GTDB taxonomy of the top 100 most represented genera, selected from the 4,125 species included in the classifier (Fig. 3). The tree was pruned from the GTDB reference tree by selecting a single representative species for each of the selected genera. Color intensity corresponds to the relative presence of flagellar proteins with higher values indicating greater abundance of that class of gene in that genus. Pie-chart indicates proportion of species in the genus that are motile (blue) or non-motile (orange) or partial protein sets (grey).

### Phylogenetic trait analysis and flagellar protein distribution

The reference bacterial tree and taxonomy files were downloaded from the Genome Taxonomy database (GTDB, release 220) to construct the phylogenetic framework for analysing flagellar protein distributions and conducting trait -based analysis. [31]. We then pruned taxa that were not in our species list using the *ape* package in R to construct a reference bacterial tree tailored to our data (Supplementary Phylogeny S1). To assess the distribution of flagellar protein homologues across bacterial genera, species were grouped into genera based on GTDB taxonomy, and a genus-level cladogram was constructed by selecting one representative species per genus from the pruned species tree (Supplementary Phylogeny S2). The 100 most represented genera in the dataset were chosen for genus-level tree (N_min =18, N_max =414). Presence-absence counts of flagellar protein homologues were aggregated at the genus level, then normalised in two steps: first, by the number of strains or species within each genus, and second, by the number of proteins in each functional class (Fig. 1C). The resulting normalised values were used to generate a heatmap, with genera arranged according to their taxonomic relationships, with the cladogram visualised in iTOL with genus names as tips. Similarly, to assess the abundance of flagellar protein homologues at the species level, presence-absence data from strains of 4,125 species present in the classifier list were aggregated at the species level. The resulting counts were normalised based on the number of strains within each species and the number of proteins in each functional class. We then used BayesTraitsV4 to model the motility trait for each species to estimate transition rates. BayesTraits is a Bayesian framework designed for model-based analyses of evolutionary selection of phenotypic or molecular traits [32]. In this approach, we analysed the trait data (predicted motility) alongside the rooted phylogenetic tree (in Nexus format) for each taxon using different models by setting transition rates to be equal and unequal for traits.

To find the best-fitting model, we used the MCMC (Markov Chain Monte Carlo); chains ran for 5 million iterations with an additional burn-in of 300,000 iterations and a sampling interval of 5,000 iterations implemented in BayesTraitsV4 [32] to estimate the log marginal likelihood for each model. The best-fitting model was determined using logarithm of Bayes Factor (logBF) to find the root probability and transition rates of different states. The mean of transition rate was calculated by taking the mean of 5,000 iterations. Ancestral character states of all nodes were estimated to identify the most likely state at each internal node in the phylogenetic tree. Histograms were generated for the distribution of transition rates using ggplot2, and posterior probabilities for each state of select nodes were tabulated using the BioGeoBEARS package in R [33]. Similarly, a presence-absence matrix was generated for each flagellar protein (1 = present, 0 = absent) across the species list as the trait data for the gain/loss estimates using Bayestraits. The multistate model then estimates the transition rates q01: The rate of protein gain (0 → 1) and q10: The rate of protein loss (1 → 0) for each ancestral node using the same parameters described above.

We then tested the correlation of the filament protein (FliC) with motility using Bayestraits. Discrete independent and dependent models were tested with the 4 states, such as 00 : non-motile-no FliC, 01 : non-motile FliC, 10 : motile - no FliC, 11 : motile-FliC. We then calculated Bayes Factors of models for each trait-fliC relationship to test whether the independent or dependent model of evolution was supported. We repeated the same analysis to test the correlation of FliC with each flagellar protein by defining the 4 possible states as follows: 00: no FliC-no Protein; 01: no FliC-Protein Presence; 10: FliC present-no Protein; 11: FliC present-Protein Presence.

## Results

We gathered 11,365 genomes from the PATRIC database and curated a search-query set of 54 flagellar proteins associated with key components of the flagellar motor from the flagellar assembly reference pathway [*map03040*] in KEGG [23] (Fig. 1A). This set includes proteins related to the C-ring, M,S,P and L rings, rod and hook, H and T rings, filament, stator, Type III secretion system, and various regulatory proteins. We then examined the presence or absence of these 54 flagellar proteins in all genomes in our dataset.

We observed a bimodal distribution of flagellar protein counts across the genomes indicating the presence of two distinct groups within the dataset. Accordingly, we assigned thresholds of 15 and 32 proteins to separate these groups (Fig. 1B). Specifically, 47% of the genomes had fewer than 15 proteins whereas 49% had more than 32. This distinct separation in flagellar protein counts allowed us to categorize the genomes into two groups: those with low counts (non-motile) and those with high counts (motile). The remaining 4% of the genomes that fall between these two groups have an intermediate number of flagellar proteins. These were classified as partial protein sets, potentially indicating an incomplete flagellar system.

We examined the functional class of flagellar proteins in both the motile and non-motile groups (Fig. 1C). The filament was found exclusively in the motile group (100% motile, 0% non-motile). Similar patterns were observed for the rod and hook (96% motile, 4% non-motile), C-ring (96% motile, 4% non-motile), MSP and L rings (94% motile, 6% non-motile), and Type-III secretion system (85% motile, 15% non-motile). For the stators, motile species still predominated (73% motile, 36% non-motile). In contrast, the proteins associated with other functional categories, such as transporters, sigma factors (59% motile, 41% non-motile), and regulators (60% motile, 40% non-motile), were more evenly distributed between the two groups. The most prevalent proteins in this group included FlrA, FliI, FlrC, RpoD, and FliY.

We investigated the distribution of flagellar proteins across the genomes though clustering and dimensionality reduction. The distance matrices calculated from the presence-absence of flagellar proteins across genomes were reduced to low dimensional embeddings using UMAP (Uniform Manifold Approximation and Projection). Clustering with DBSCAN identified three major clusters, with some of which contain subclusters that exhibit distinct compositions (Fig. 1D; Supplementary Fig.S2). These subclusters represent finer divisions within the major clusters. Cluster I comprised 47.1% of the genomes and consisted predominantly of non-motile genomes from the most abundant phyla, including *Actinomycetota, Pseudomonadota, Bacillota, Bacteroidota*, and *Cyanobacteriota*. Subclusters II, III, IV, and V together formed a major cluster, with Subclusters II, III, and IV accounting for 27.7%, 15.2%, and 6.3% of the genomes, respectively, and classified as motile. Subclusters III and V were primarily composed of genomes from the phylum *Pseudomonadota*, whereas Cluster IV contained genomes from *Bacillota*. Subcluster V and Cluster VI contained genomes classified as partial protein sets, with Cluster VI primarily consisting of genomes from the phylum *Chlamydiota*. The genomes classified as partial protein sets were nearer to the motile cluster than the non-motile cluster.

We then validated our classifier based on the counts of flagellar proteins using published phenotypic data [13, 21, 34]. Observations of motility for Firmicutes curated from Bergey’s Manual of Systematic Bacteriology[34] demonstrated an accuracy rate of 95%, with 159 accurate predictions and nine mismatches (Supplementary Fig. S3). We further compared phenotypic data from 1937 species curated in a previous study [21] and accurately classified 808 species as motile and 867 as non-motile with an overall accuracy of 86.5% (Supplementary Fig. S4).

We examined the proportion of flagellar proteins in each class for the most prevalent genera in our dataset (Fig. 2). Species from the phyla *Pseudomonadota, Campylobacterota*, and *Spirochaetota* showed a consistently high abundance of flagellar components, with nearly complete set of structural and regulatory proteins. In contrast, members from the phyla *Bacteroidota, Bacillota*, and *Actinomycetota* were missing proteins in the filament, C-ring and MSP-& L-ring clusters. Type-III Secretion System (T3SS) proteins and stator proteins were present in all genera (Supplementary Fig. S5). In particular, FliI, an ATPase[35, 36] in the T3SS, and FliA, a sigma factor[37, 38], were conserved across all genera (Fig. 2).

We measured the proportion of motile and non-motile genomes containing filament, stator, and T3SS proteins across all motile and non-motile proteomes (Supplementary Fig. S6). We observed 100% of motile proteomes had at least one filament hit, and 35% of motile genomes contained a hit for all four query proteins. Alternatively, ∼99% of the non-motile genomes showed no hit for any filament query protein, suggesting a strong correlation of filament proteins with motility. The full set of T3SS proteins was found in 21% of motile genomes and 57% of motile genomes had all T3SS proteins except FlhE. Conversely, 97% non-motile genomes had either FliH and FliI (57%) or FliI alone (40%). The split across motile and non-motile genomes for the stator proteins was more nuanced, with FliA, MotA, and MotB displaying the highest proportion of hits. FliA was hit in 41% of non-motile genomes, and FliA, MotA, and MotB were hit altogether in 20% on non-motile genomes.

We employed trait analysis using BayesTraits [32] to infer the ancestral states and transition rates between motile and non-motile across a pruned phylogenetic tree comprising 4,125 bacterial species from GTDB matching our species list (Fig. 3). Among the models tested, the multistate model with qMN ≠ qNM rates provided the best fit (logBF = 76.4) for our data, Supplementary Table S4). The multistate model yielded a mean root probability of 0.99931 for the motile trait and 0.00069 for the non-motile trait. Across the phylogeny, transitions from motile to non-motile states (qMN) were more frequent, with a mean transition rate of 0.8058, whereas transitions from non-motile to motile (qNM) had a lower transition rate of 0.21987 (Supplementary Fig. S7). We observed that 54% of the internal nodes exhibited high probabily for the motile state, whereas 45% of the nodes (predominantly Actinomycetota, Bacteroidota, Pseudomonadota, Bacillota, and Cyanobacteriota) displayed high probabilities for the non-motile state, suggesting shifts toward non-motility in these lineages. We further employed trait analysis for MotA, MotB, FliC, FliH, and FliI for comparison of clading across the same tree (Supplementary Fig. S8)

**Fig. 3:**
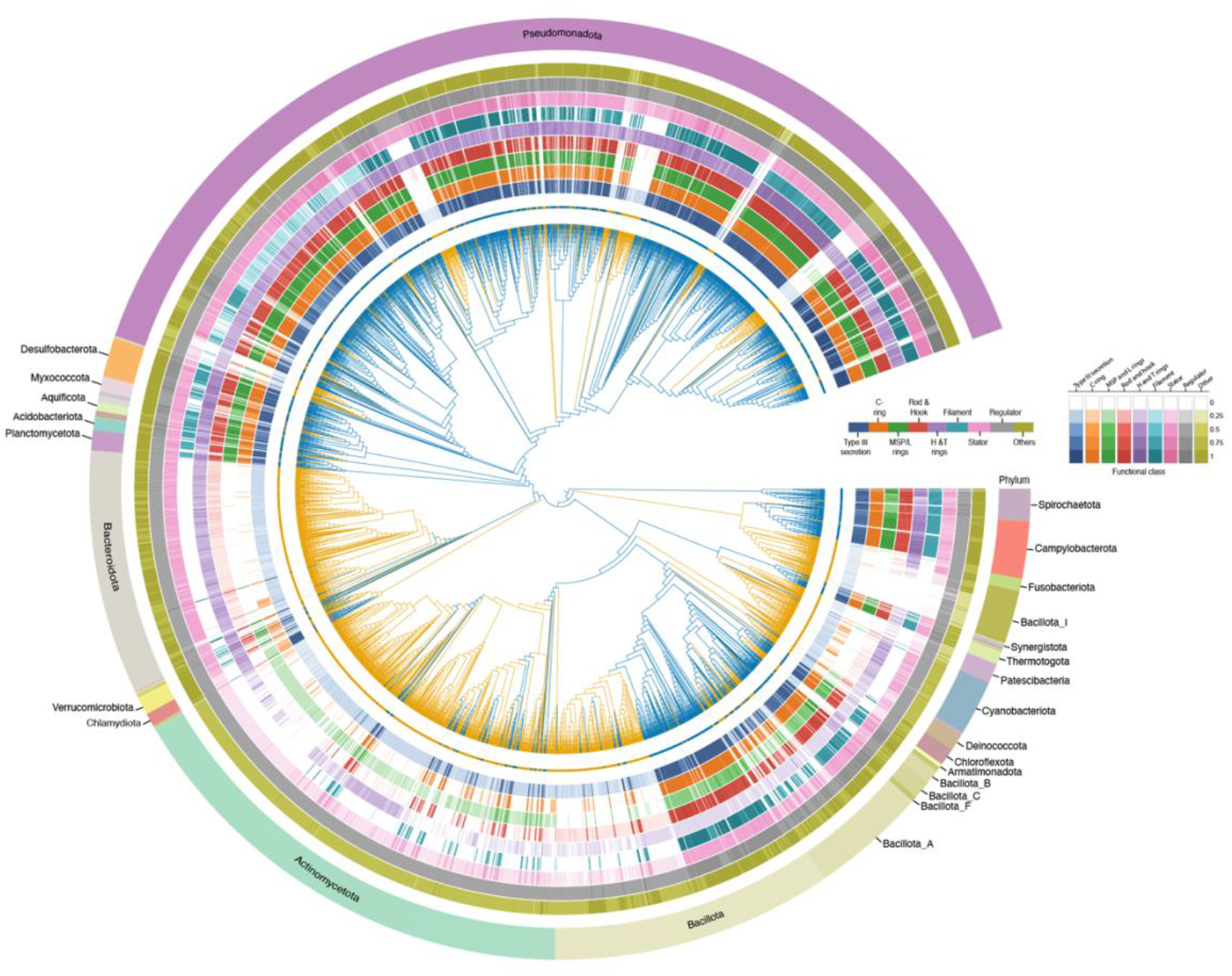
Bayesian reconstruction of ancestral motility traits. Phylogenetic tree depicting the relationships among 4,125 bacterial species common to our dataset and the Genome Taxonomy Database (GTDB). Tree is pruned from the GTDB reference tree for the matching 4125 bacterial species. Branch colours represent the posterior probabilities (Pmot(θ| X)) of the motility states estimated for each node using the Markov Chain Monte Carlo (MCMC) multistate model in BayesTraits. Branch colours indicate the following states: orange for non-motile (Pmot(θ| X)>0.5), blue for motile (Pmot(θ| X)<0.5), and grey for nodes having equal probabilities for both traits (Pmot(θ| X)=0.5). The nine inner rings are color-coded according to functional class (blue: T3SS; orange: C-ring; green: MSP/L rings; purple: H and T rings; cyan: Filament; pink: Stator; grey: Regulator; olive: Other), while the outer ring is color-coded by phylum. For each functional class, circular heatmaps depict the normalised abundance of flagellar protein homologues at the species level, with presence-absence counts aggregated at the species level and normalised by both the number of strains and proteins within each functional class (Fig. 1C).

We measured the gain and loss rates for each protein and the correlation of each protein with motility (Supplementary Fig. S9, Supplementary Table S5). All proteins had significantly different rates of transition for gain and loss (logBF > 2) except MotB, FliI, MotY, FlgD and RpoN. Proteins with different transition rates for gain and loss, including all filament, rotor, secretion, and rod and hook proteins, had higher loss than gain (qMN < qNM). The three exceptions, which had higher gain than loss (qMN < qNM), were MotA, FliA, and MotX. We confirmed that FliC was highly correlated with motility (logBF = 1993.88) and then measured the correlation of each protein with FliC (Supplementary Table S6). For the non-filament proteins, the most correlated with FliC (logBF > 1000), and accordingly with motility, were FliEFGKLMNOPQR, FlhAB, and FlgBCDEGHIKLN. All filament query proteins were strongly correlated with FliC (logBF > 300). We then compared the gain and loss of each protein with respect to FliC (Supplementary Fig. S10). Proteins with a higher rate of gain that was at least tenfold greater than the gain rate of FliC included FliA, FliH, MotX, and FlgD, and those with a lower rate of gain as least tenfold lower than that of FliC included FliI, FliT, MotC, FlhD, FlhE, FlgH, FlgI, and FlgQ. Similarly, proteins with a higher rate of loss that was at least tenfold greater than that of FliC included FliT, MotD, FlhC, FlhE, and FlgO, and with a lower rate of loss as least tenfold lower than that of FliC included FliI.

## Discussion

### Classifier for motility

In this work we developed a classification system that can accurately predict whether a taxon is motile by quantifying the presence and absence of flagellar proteins in a given genome. We validated this system across a large dataset of motility data using published datasets and Bergey’s Manual of Systematic Bacteriology [34]. Our system performs well: it is 95% accurate across *Firmicutes* (Supplementary Fig. S3) using annotations in Bergey’s Manual of Systematic Bacteriology [13, 34] and displays 87% sensitivity (true-positive rate) and 86% specificity (true-negative rate) across all prokaryotes using curated metadata from a previous study [21] (Supplementary Fig. S4).

### Common bacterial ancestor was motile

We confirmed, as reported elsewhere [12], that the last bacterial common ancestor (LBCA) likely contained a flagellar motor (posterior probability of 0.99). Our approach uses a combination of methods combining sequence homology (Jackhmmer), structural homology (Foldseek, AlphaFold), dimensionality reduction (UMAP, PCA) and large well-curated databases (KEGG, PATRIC). We reconstructed motility across the phylogeny to show that motility was primarily lost in a few phyla (predominantly *Actinomycetota, Bacteroidota, Pseudomonadota, Bacillota*, and *Cyanobacteriota*).

### Filament proteins are informative in simple motility assessment

By far the best, and most simple determinant for flagellar motility is the presence or absence of a filament. Considering only motile genomes, 49% contained hits for all four filament proteins in our query set (FliC, FliD, FliS, FliT), and 99.75% contained at least one filament hit. If our classification system were reduced to a search only for FliC, it would still be 99.5% accurate (Supplementary Fig. S11, Supplementary Fig. S12). Overall, a search for FliC is a good, coarse assessment of whether a species is motile or not.

### What use is half a motor?

Cases where there were some, but not all, of the flagellar proteins are worthy of further consideration. To paraphrase Mivart ‘what use is half a motor?’ [39, 40]. Our model, and experimental experience[41, 42], asserts that an incomplete flagellar system cannot confer motility, and we validate this, but the question remains how did a half motor come to be? If an essential rotor protein such as FliG was lost, are the other proteins only remnants of a motor long since lost? By and large these loss events were long ago. Non functional remnants of flagellar proteins have been observed in several species [43, 44]. We observed that <0.3% (34/11365) of all species had at least one filament protein but no rotor proteins. However, 4% of motile species had a filament but less than a full rotor (ie FliGMN). This raises the question of what is the minimal rotor that can sustain motility. There are species such as *Aquifex aeolicus* which have been observed to be missing FliM [45], however in our search we did observe a homolog for FliM in *Aquifex aeolicus* (GenBank AAC07248.1, RefSeq assembly GCF_000008625.1). Conversely, we saw that 11% of non-motile species had at least one rotor protein (FliG, FliM, FliN) but no filament proteins. Whilst we would expect to find a high number of T3SS and stator homologues in non-motile genomes as these are well known to have non-flagellar homologues [46–50] the presence of rotor proteins without a filament raises the possibility that rotor components could have an alternative function.

Incomplete modules of the motor may also represent remnants of a motor following gene loss. *Rhodobacter sphaeroides* possess two flagellar systems, with one acquired through HGT from a γ-proteobacterium, demonstrating that entire flagellar gene sets can be transferred between species across large phylogenetic distances [51, 52]. It is plausible that key flagellar genes could be lost to render a species non-motile. Filament proteins like FliC make up the vast mass of the flagella and demand a high energy cost for synthesis [53, 54]. If a bacterium invests such energy to make a filament, it requires a motor and stator complex to make that cost worthwhile and confer a selective advantage. However, if the filament gene is lost in a loss event, then the energy cost to maintain, for example, a few rotor proteins, is comparatively low.

### Exceptions to the rule

We can generate hypotheses surrounding the few strains that appear misclassified using published datasets [13, 21] (≤13%, Supplementary. Fig S3, Supplementary. Fig S4, Supplementary Table S7). In the case of the *Firmicutes* data set [13], there were 8 exceptions, with 7 species reported as non motile that were classified as motile and one species reported as motile that was classified as non-motile (Supplementary Fig. S3). For the wider dataset of 1937 species [21], 119 were incorrectly classified as motile and 143 incorrectly classified as non-motile. We considered experimentally testing specific strains to execute our own single-cell motility tests, however many of these strains are hard to obtain and it is uncertain whether they would be amenable to lab-based motility assays which may require specific environmental stimuli to express motility genes [55]. Experimental testing of these outliers, such as by culturing and directly testing for and measuring motility [56, 57] could further validate and refine the accuracy of our classifier, and identify where phenotypic descriptions compiled in metastudies contained errors.

### Comparing and correlating protein loss across all flagellar proteins

The majority of flagellar proteins (87%) had higher loss rates than gain rates demonstrating that flagellar motility is more easily lost than gained. We calculated the presence and absence of FliC along with motility traits and reconstructed it across our reference GTDB tree of 4125 species. We found strong evidence for a dependent model of trait evolution (logBF = 1993.88) (Supplementary Table S6) indicating a close interconnection between flagellar motility and the presence/absence of the FliC. This further shows the utility of searching for homologues of FliC as a quick assay for flagellar motility when exploring a new genome. Measuring correlation between each protein with FliC allows us to assert a hierarchy of the most important proteins for motility. Proteins in the MS- and C-rings – the rotor – are highly correlated with FliC, as are Rod, PL-rings and hook. This is logical, these are essential components that couple rotation to the filament and assembly of the filament depends on their presence [58]. MotA and MotB are highly correlated with each other (logBF_MotA-MotB_ = 622), comparable with their separate correlation with FliC (logBP_MotA-FliC_ = 655; logBP_MotB-FliC_ = 884). Ancestral state reconstruction (ASR) across our bacterial phylogeny indicates that MotA and MotB have largely been lost in the same clades as FliC. (Supplementary Fig. S8). In contrast, conserved ATPase proteins of the T3SS [35], FliH and FliI, are present in both the *Actinomycetota* and *Bacteroidota* clades, whereas FliC is not, and FliI is present in nearly all (99.6%) of our proteomes. Overall, this validates our trait analysis of motility across the tree of life as well-known, conserved flagellar proteins display the same shared phylogenetic history.

### Conclusions

Given the strong correlation of FliC with motility, the high energetic cost of filament assembly [58] and the benefit to host-associated pathogens to lose the filament [59], it is tempting to speculate that loss of motility begins with loss of FliC. However, we cannot ascertain the order of loss events and losing other proteins that assemble before the filament would also destroy filament formation [60].

Correlation with other traits (such as correlating loss of motility in the *Bacteroidota* with the gain of gliding motility via *gldKLMN*[61], remains to be tested across a larger reference tree. It may be interesting to consider exactly when additional flagellar genes such as those present only in *Campylobacterota* arose [60, 62, 63]. These loss and gain events are likely in the distant past and accordingly analyses will be subject to substantial noise. Overall, better reconciliation of gene trees with reference phylogenies, combined with ecological data, will help to quantify ecological shifts in flagellar motility throughout history.

## Supporting information

Supplementary Phylogenies 1&2, Supp. Table S1-3,S7

## Data availability

All the data used is presented as tables in the Supplementary Information

## Code availability

The code used is available at https://github.com/BakerLabBABS/classifier

## Author information

### Contributions

JSP and MABB designed the analysis. JSP wrote original code, executed all final analysis including search, identification, classification and trait analysis. SG performed initial survey of genes in PATRIC database. JS examined phenotypes and provided validation of classifier. CPL assisted with structural homology. NJM and LM assisted with trait analysis and tree visualisation. MAAB. supervised the design, execution, and writing of the project. All authors contributed to writing and revision of the manuscript.

## Acknowledgements

Funding: MABB is supported by US Navy Office of Naval Research, Research grant no. N62909-22-1-2051. MABB, LM and NJM were supported by HFSP Project Grant RGY0072/21. MABB and NJM are supported by Australian Research Project Discovery Project DP240100462.

## Supplementary Information

**Supplementary Fig. S1**. DBSCAN clustering of UMAP embeddings for (A) FliI and (B) FliG. Pairwise distance matrices from multiple sequence alignments were reduced to low-dimensional embeddings using UMAP and clustered with DBSCAN. Cluster annotations were assigned based on information retrieved from UniProt using ID mapping between PATRIC genome IDs and UniProt IDs. Each point represents a protein sequence, with colours indicating different clusters.

**Supplementary Fig. S2**. DBSCAN clustering of UMAP embeddings for the motility classifier. The distance matrices based on the presence-absence of flagellar proteins across genomes were reduced to low-dimensional embeddings using UMAP, and then clustered using the DBSCAN.

**Supplementary Fig. S3. Confusion Matrix of Motility Classifications**. Matrix comparing the motility classifications of 169 species from our classifier dataset with those from the Firmicutes motility dataset by Fouad El Baidouri et al. (2016).

**Supplementary Fig. S4. Confusion Matrix of Motility Classifications**. Matrix comparing the motility classifications of 1937 species in our classifier dataset with those from the curated dataset from a previous study [21].

**Supplementary Fig. S5**. Heatmap showing the the normalized abundance of flagellar protein homologues, specifically filament, Type-III Secretion System (T3SS), and stator proteins, across the most prevalent genera (Fig. 2). Data were derived from species with available GTDB lineage information. Presence-absence counts were aggregated at the genus level and normalized based on both the number of strains within each genus and the number of proteins in each functional class (Fig. 1C). The adjacent cladogram illustrates the GTDB taxonomy of the top 100 most represented genera, selected from the 4,125 species included in the classifier (Fig. 3). The tree was pruned from the GTDB-reference tree by selecting a single representative species for each of the selected genera. Color intensity corresponds to the relative presence of flagellar proteins with higher values indicating greater abundance in that genus.

**Supplementary Fig.S6**. Sankey plot showing the distinct patterns of flagellar proteins across motile and non-motile genomes for each flagellar component: A) Filament, B) Stator, and C) Type III secretion system (T3SS), D) Rotor. In each panel, genomes are first divided into *motile* and *non-motile* categories. Subsequent branches display individual proteins/different combinations associated with each component of the flagellar apparatus. *Other combinations* indicate under-represented combinations not shown individually, whereas *None* represents genomes lacking those proteins. The width of each flow is proportional to the number of genomes.

**Supplementary Fig. S7. Transition rates of motility**. Histograms showing the posterior distribution of transition rate estimates between motile (qMN) and non-motile (qNM). The red dotted line indicates the mean, and the blue line indicates the 95% CI.

**Supplementary Fig.S8:** Bayesian ancestral state reconstruction of the gain and loss of flagellar proteins across the bacterial phylogeny: (A) FliH, (B) FliC, (C) FliI, (D) MotA, and (E) MotB. The phylogenetic tree depicts the relationships among 4,125 bacterial species common to our dataset and the Genome Taxonomy Database (GTDB). This tree is pruned from the GTDB reference tree for the matching 4125 bacterial species. Branches are colored according to the posterior probability of protein presence (gain) or absence (loss) at each node, as estimated using the Markov Chain Monte Carlo (MCMC) multistate model in Bayestraits. Blue indicates a higher probability of protein presence whereas orange indicates a higher probability of protein absence.

**Supplementary Fig. S9:** Bar Plot showing the log-transformed gain/loss ratio (ln(q01/q10)) for each flagellar protein, with colors indicating their associated flagellar substructures. Positive values indicate a higher frequency of gain events, whereas negative values indicate a higher frequency of loss events. Values close to zero indicate similar rates of gain and loss.

**Supplementary Fig. S10:** Bar plot showing the log-transformed normalized gain and loss rates for each protein relative to FliC. The y-axis represents **ln (q**_**01**_ **/ q**_**01, FliC**_) where q_01_gain_ and q_10_loss_ indicate the gain or loss rates of a given protein, normalized against the respective rates for FliC. A **Positive value** indicates a higher gain rate compared to FliC and **negative value** indicates a lower gain rate compared to FliC whereas **zero** indicates same gain rate as FliC. Bars are grouped by proteins within different substructures, with separate bars for gain and loss rates.

**Supplementary Fig. S11: Confusion Matrix of Filament Classifications**. This matrix quantifies presence and absence of filament proteins (FliC, FliD, FliS, FliT) for motile and non-motile clusters for 10960 species present in our dataset (full dataset with those genomes classified as partial removed). Accuracy is calculated at 99.6% as: Accuracy = (TP + TN) / (TP + TN + FP + FN)

**Supplementary Fig. S12: Confusion Matrix of FliC *Classification***. This matrix quantifies presence and absence of FliC or motile and non-motile clusters for 10960 species present in our dataset (full dataset with those genomes classified as partial removed). Accuracy is calculated at 99.5% as: Accuracy = (TP + TN) / (TP + TN + FP + FN)

**Supplementary Table S1**: List of species and their corresponding PATRIC genome IDs used in this study.

**Supplementary Table S2:** The complete list of species and flagellar proteins, together with the total number of proteins present.

**Supplementary Table S3:** Table showing the percentage of sequences removed for each flagellar protein during the length-filtering step.

**Supplementary Table S4**: Log Marginal Likelihood (log ML) estimates using the BayesTraits Multistate algorithms for equal and unequal rates for qMN and qNM with MCMC and a stepping stone sampler for motility traits.

**Supplementary Table S5:** Log Marginal Likelihood (log ML) estimates using the BayesTraits Multistate algorithms for equal and unequal rates for q01(gain) and q10(loss) with MCMC and a stepping stone sampler for individual flagellar proteins.

**Supplementary Table S6:** Log Marginal Likelihood (log ML) estimates using the BayesTraits Multistate algorithms for dependent and independent models for correlated evolution of other proteins vs FliC.

**Supplementary Table S7:** The list of species with mismatched motility classification as compared with the phenotypic data curated from published literature [13, 21].

**Supplementary Phylogeny S1:** Phylogenetic tree of 4125 species pruned from reference GTDB tree in Nexus format.

Supplementary Phylogeny S2: Phylogenetic tree of 100 representative genera pruned from reference GTDB tree using one species per genus in Newick format.

