## Supplementary Figs S1-S12, Supplementary Table S4-6. for "Mapping the loss of flagellar motility across the tree of life"

### SUPPLEMENTARY FIGURES

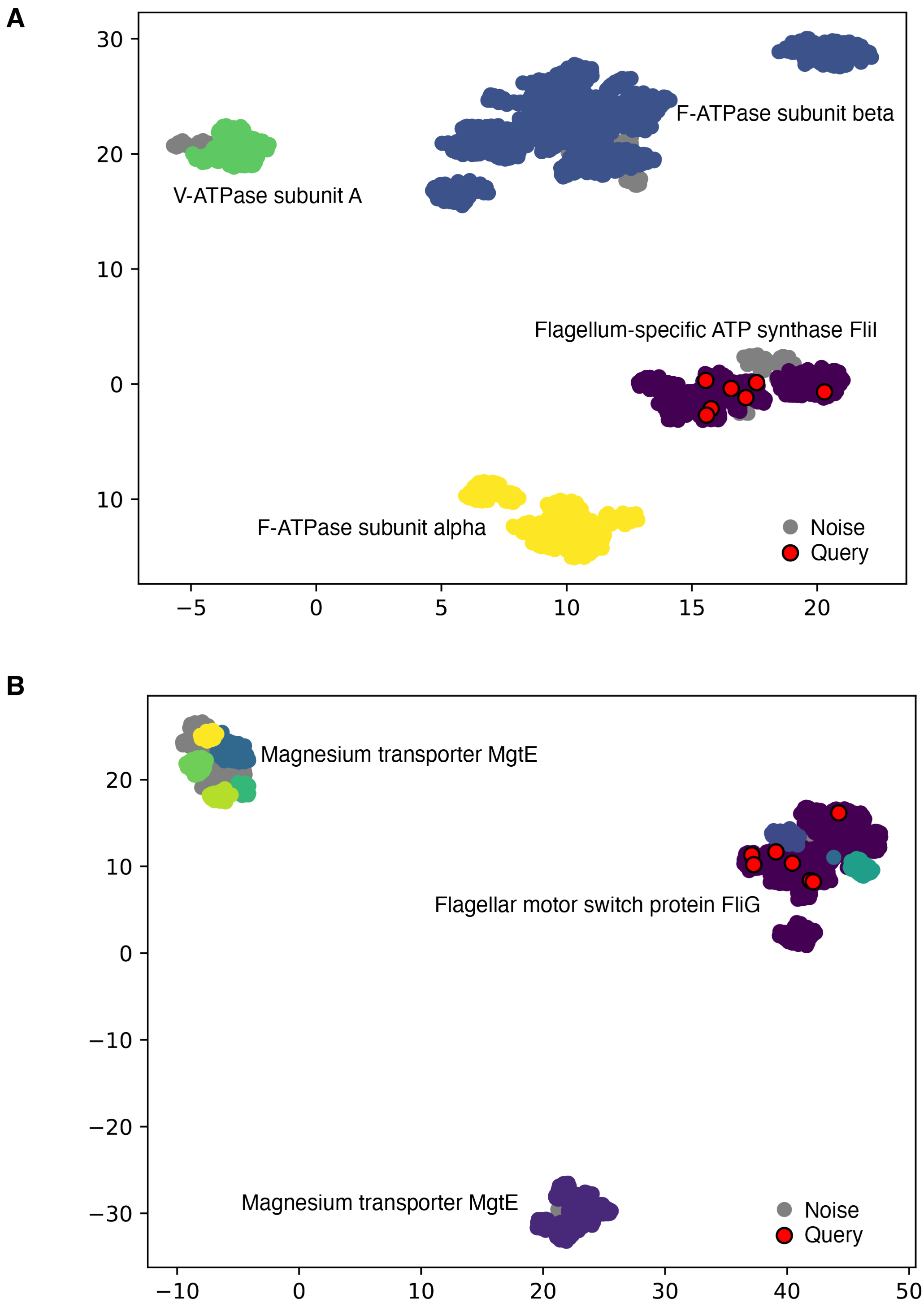

**Supplementary Fig. S1.** DBSCAN clustering of UMAP embeddings for (A) FliI and (B) FliG. Pairwise distance matrices from multiple sequence alignments were reduced to low-dimensional embeddings using UMAP and clustered with DBSCAN. Cluster annotations were assigned based on information retrieved from UniProt using ID mapping between PATRIC genome IDs and UniProt IDs. Each point represents a protein sequence, with colours indicating different clusters.

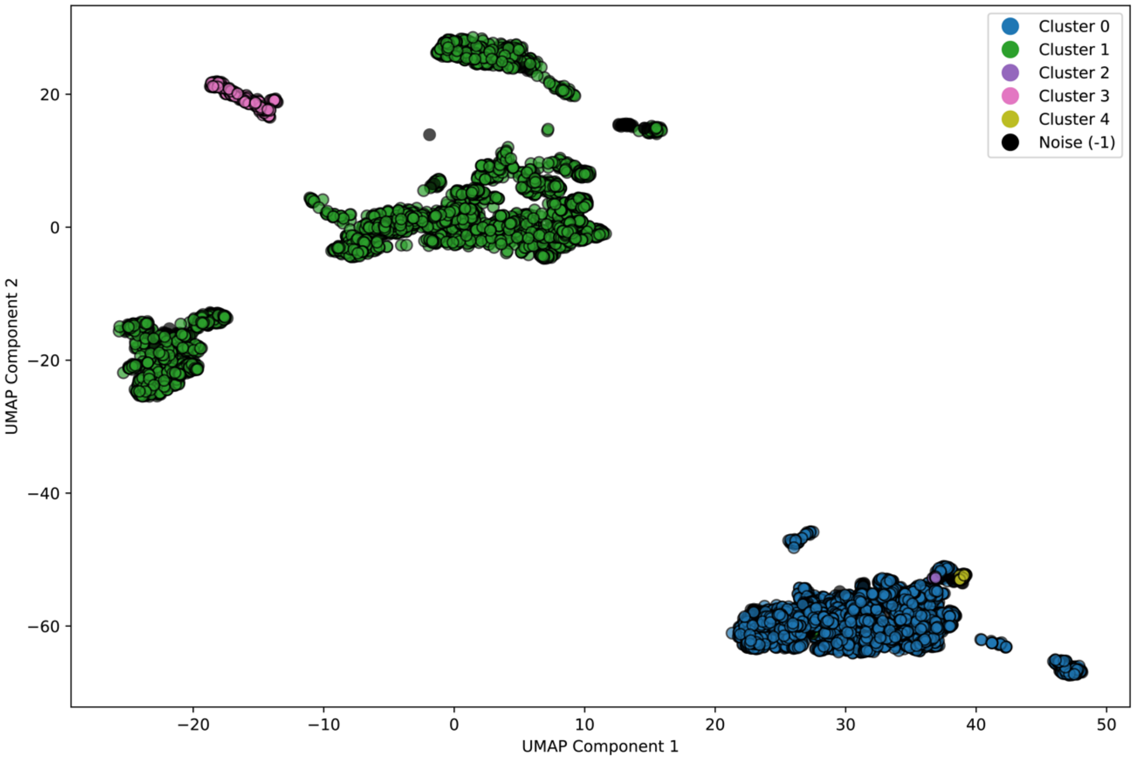

**Supplementary Fig. S2.** DBSCAN clustering of UMAP embeddings for the motility classifier. The distance matrices based on the presence-absence of flagellar proteins across genomes were reduced to low- dimensional embeddings using UMAP, and then clustered using the DBSCAN.

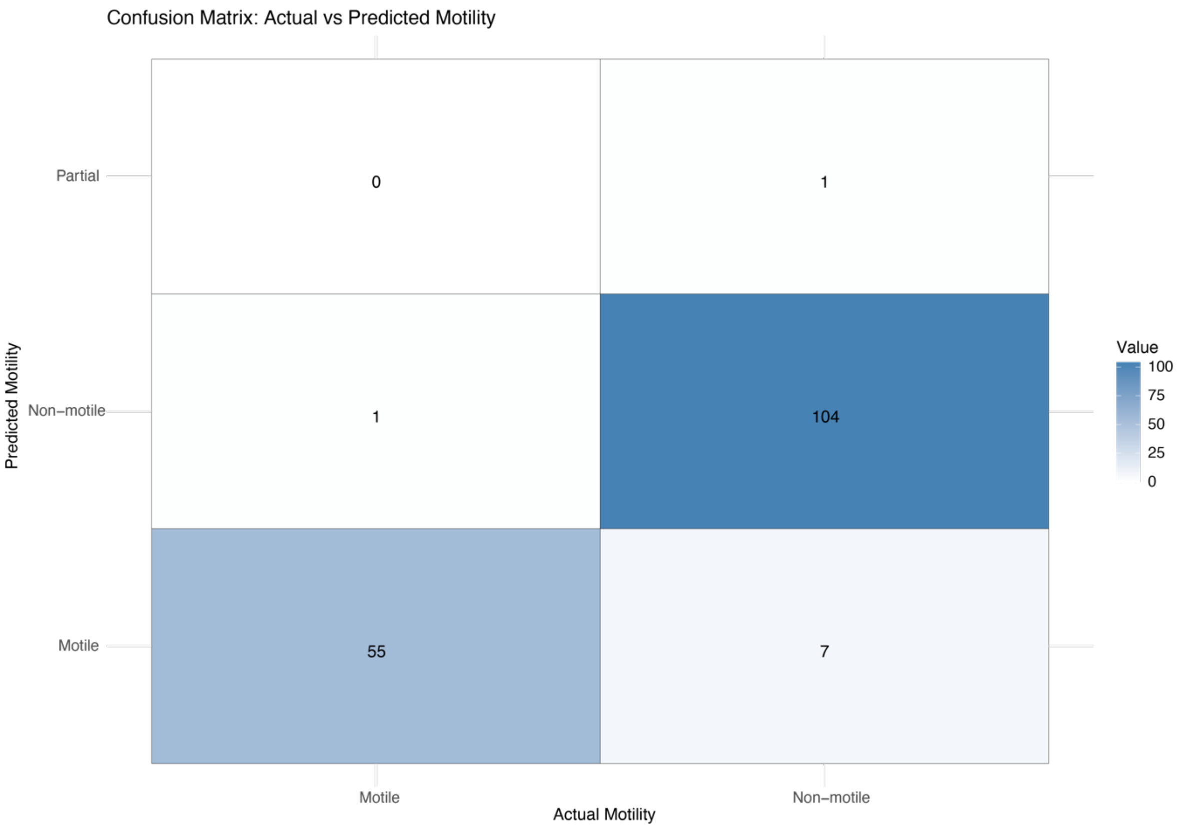

**Supplementary Fig. S3. Confusion Matrix of Motility Classifications.** This confusion matrix compares the motility classifications of 169 species from our classifier dataset with those from the Firmicutes motility dataset by Fouad El Baidouri et al. (2016).

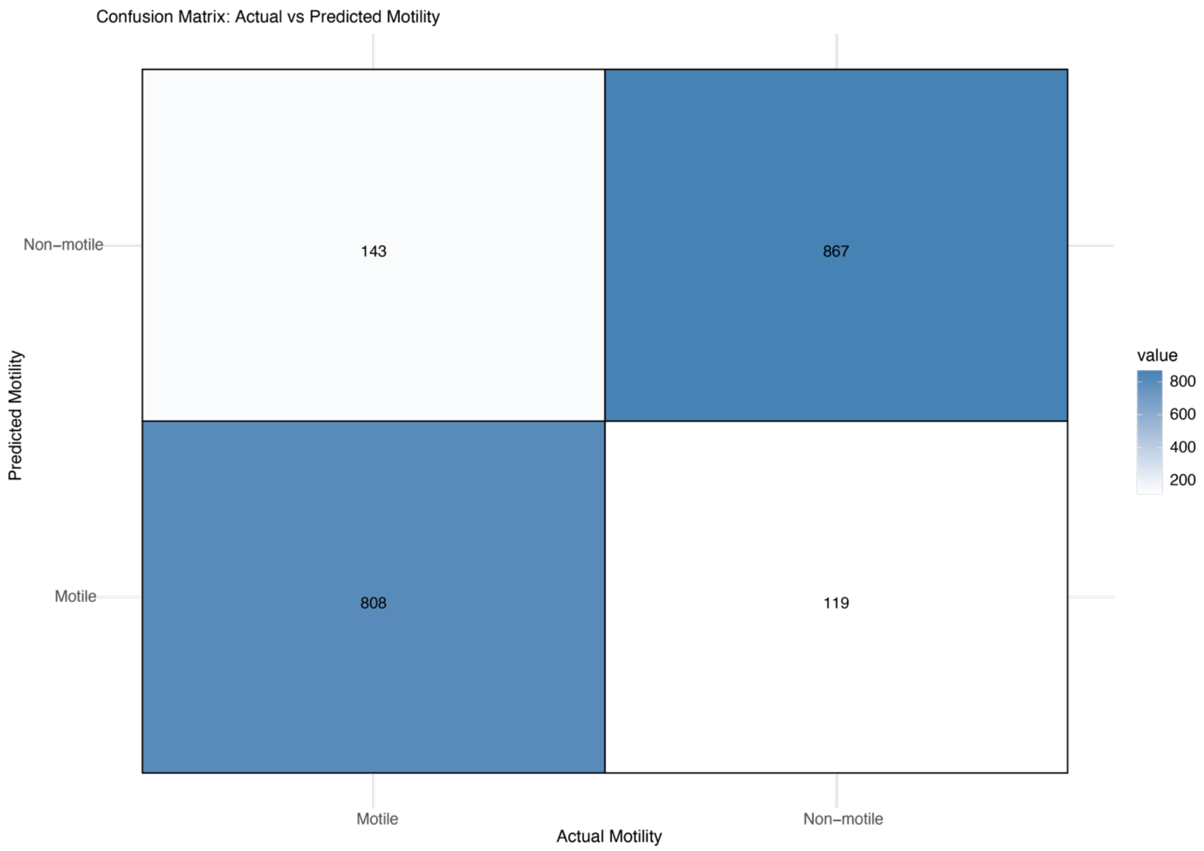

**Supplementary Fig. S4. Confusion Matrix of Motility Classifications.** This matrix compares the motility classifications of 1937 species in our classifier dataset with those from a previous study (21).

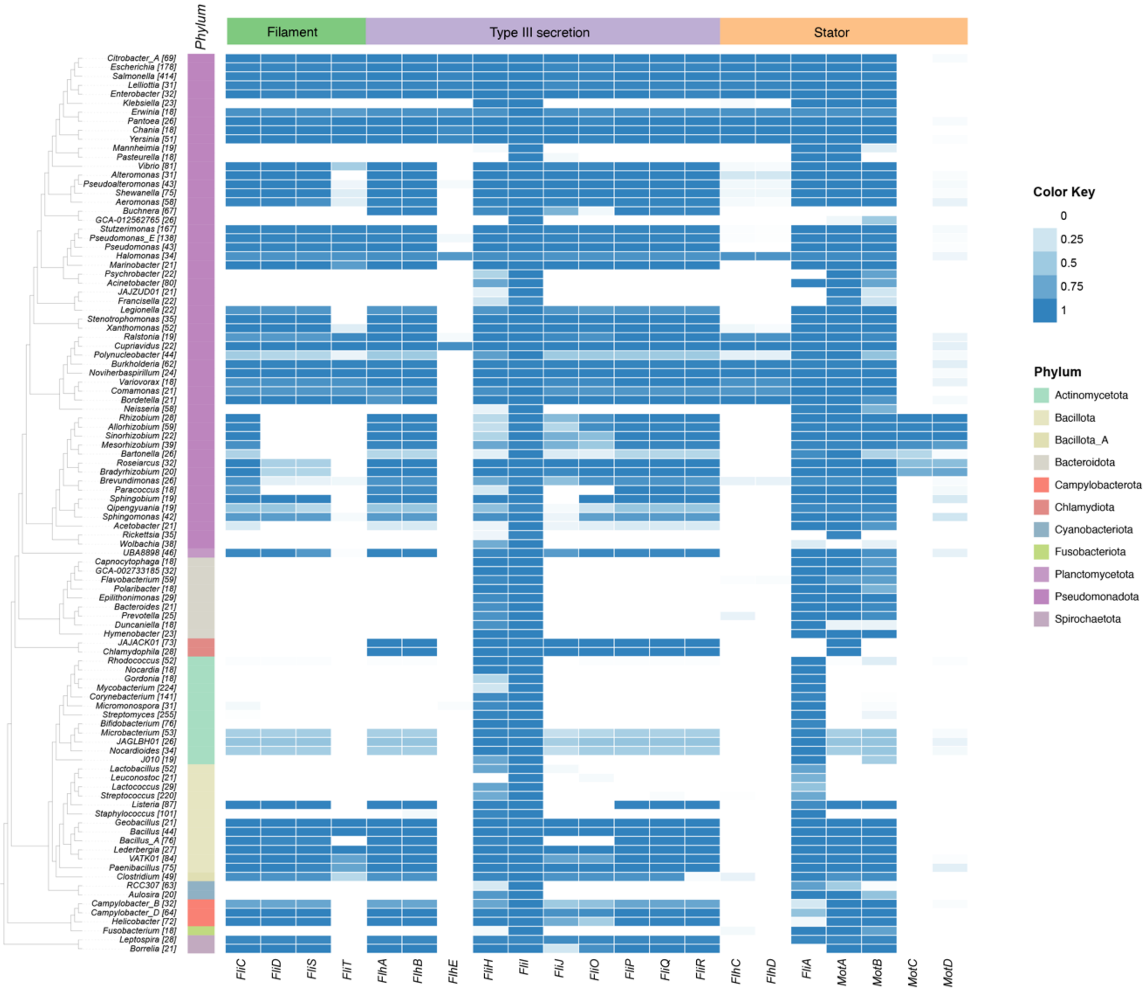

**Supplementary Fig. S5.** Heatmap shows the detailed breakdown of the normalized abundance of flagellar protein homologues, specifically filament, Type-III Secretion System (T3SS), and stator proteins, across the most prevalent genera (Fig. 2). Data were derived from species with available GTDB lineage information. Presence-absence counts were aggregated at the genus level and normalized based on both the number of strains within each genus and the number of proteins in each functional class (Fig. 1C). The adjacent cladogram illustrates the GTDB taxonomy of the top 100 most represented genera, selected from the 4,125 species included in the classifier (Fig. 3). The tree was pruned from the GTDB-reference tree by selecting a single representative species for each of the selected genera. Color intensity corresponds to the relative presence of flagellar proteins with higher values indicating greater abundance in that genus.

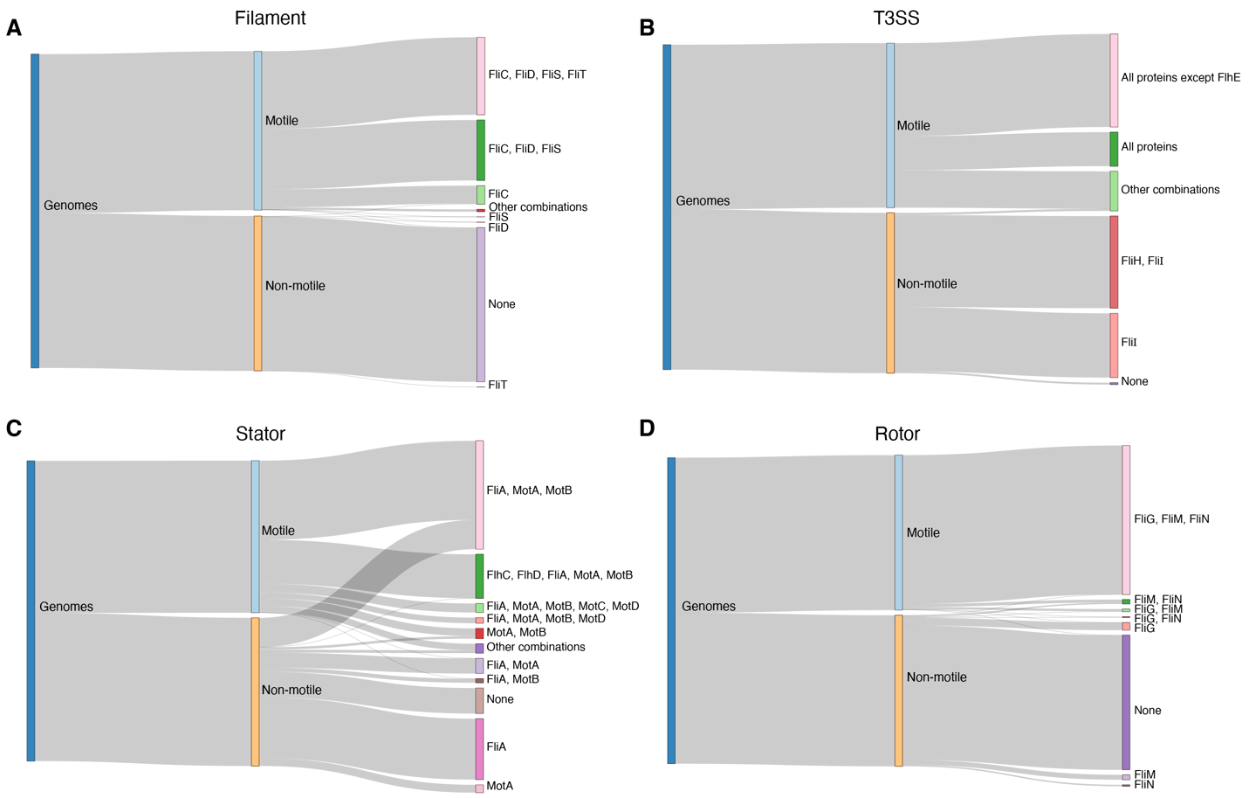

**Supplementary Fig.S6.** Sankey plot showing the distinct patterns of flagellar proteins across motile and non-motile genomes for each flagellar component: A) Filament, B) Stator, and C) Type III secretion system (T3SS), D) Rotor. In each panel, genomes are first divided into *motile* and *non-motile* categories. Subsequent branches display individual proteins/different combinations associated with each component of the flagellar apparatus. *Other combinations* indicate under-represented combinations not shown individually, whereas *None* represents genomes lacking those proteins. The width of each flow is proportional to the number of genomes.

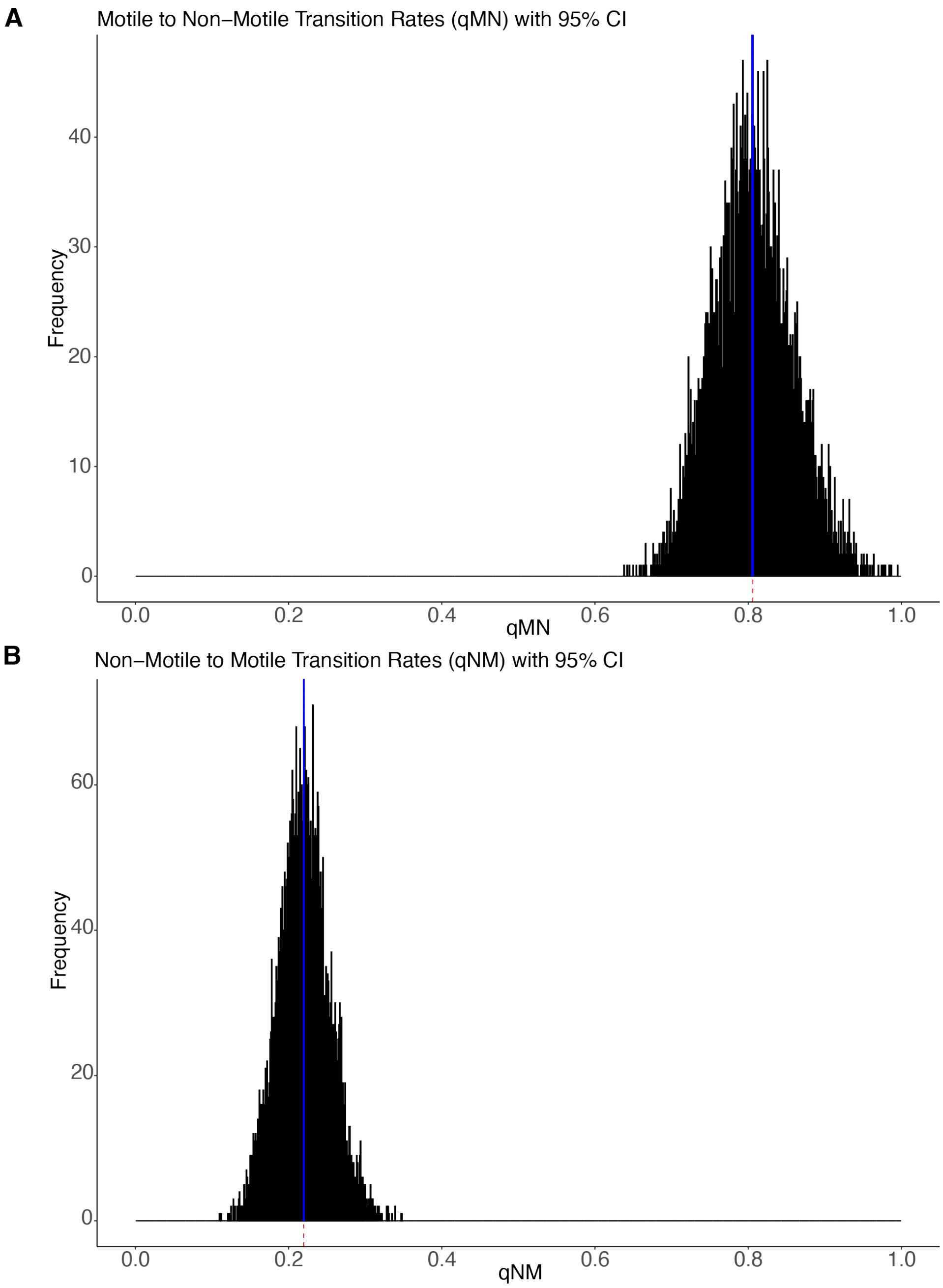

**Supplementary Fig.S7. Transition rates of motility.** Histograms indicate the posterior distribution of transition rate estimates between motile (qMN) and non-motile qNM). The red dotted line indicates the mean, and the blue line indicates the 95% CI.

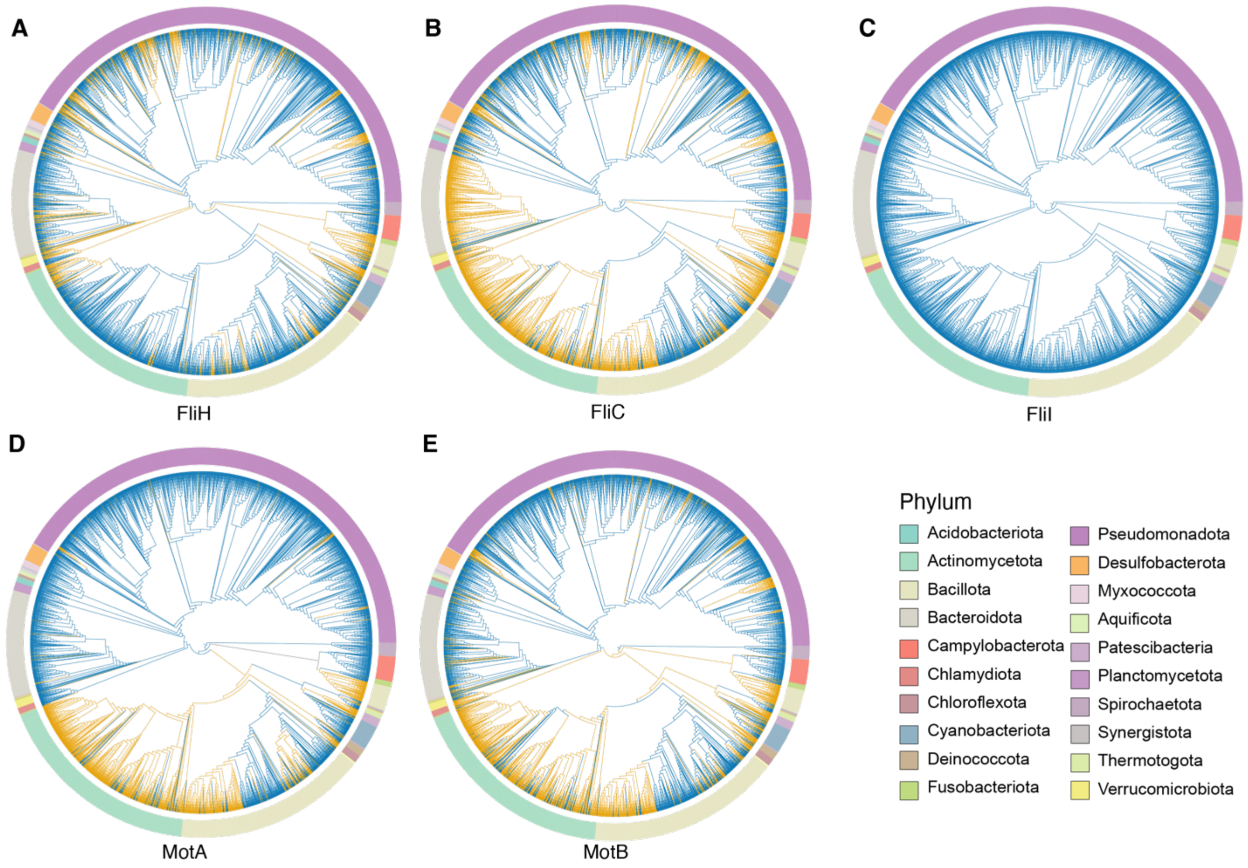

**Supplementary Fig.S8:** Bayesian ancestral state reconstruction of the gain and loss of flagellar proteins across the bacterial phylogeny: (A) FliH, (B) FliC, (C) FliI, (D) MotA, and (E) MotB. The phylogenetic tree depicts the relationships among 4,125 bacterial species common to our dataset and the Genome Taxonomy Database (GTDB). This tree is pruned from the GTDB reference tree for the matching 4125 bacterial species. Branches are colored according to the posterior probability of protein presence (gain) or absence (loss) at each node, as estimated using the Markov Chain Monte Carlo (MCMC) multistate model in Bayestraits. Blue indicates a higher probability of protein presence whereas orange indicates a higher probability of protein absence.

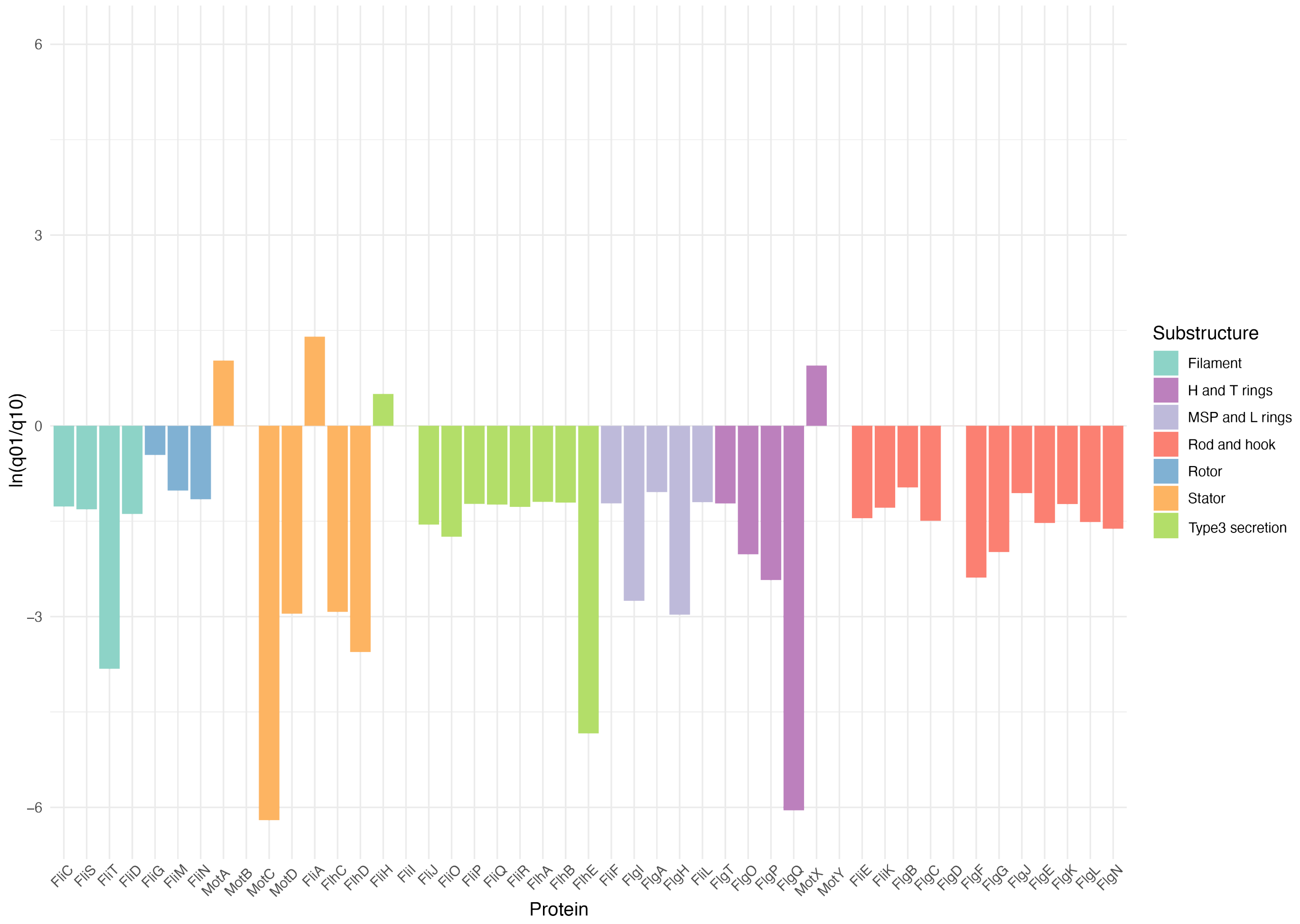

**Supplementary Fig. S9:** Bar Plot showing the log-transformed gain/loss ratio (ln(q01/q10)) for each flagellar protein, with colors indicating their associated flagellar substructures. Positive values indicate a higher frequency of gain events, whereas negative values indicate a higher frequency of loss events. Values close to zero indicate similar rates of gain and loss.

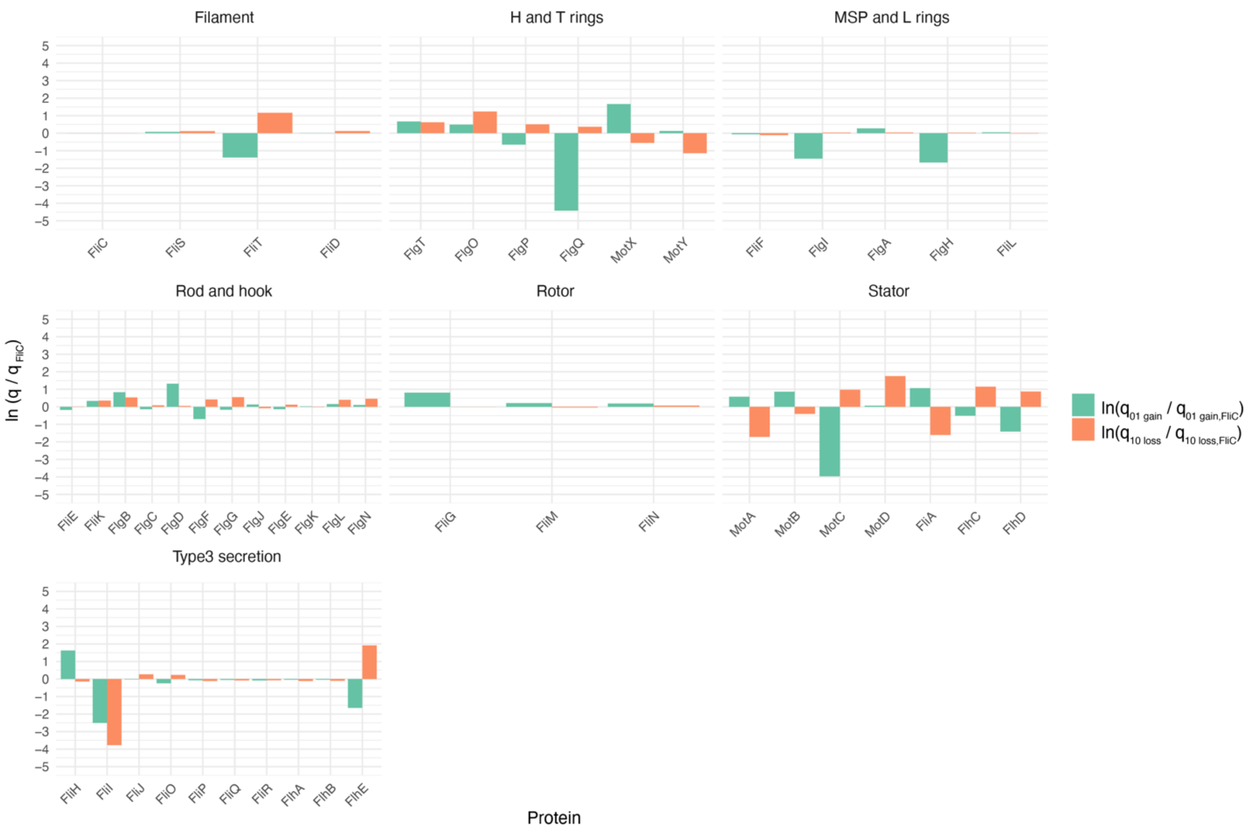

**Supplementary Fig. S10:** Bar Plot showing the log-transformed normalized gain and loss rates for each protein relative to FliC. The y-axis represents **ln ($q_{01\_gain}$ / $q_{01\_gain,\ FliC}$) or ($q_{10\_loss}$ / $q_{10\_loss,\ FliC}$))**, where $q_{01\_gain}$ and $q_{10\_loss}$ indicate the gain or loss rates of a given protein, normalized against the respective rates for FliC. A **Positive value** indicates a higher gain rate compared to FliC and **negative value** indicates a lower gain rate compared to FliC whereas **zero** indicates same gain rate as FliC. Bars are grouped by proteins within different substructures, with separate bars for gain and loss rates.

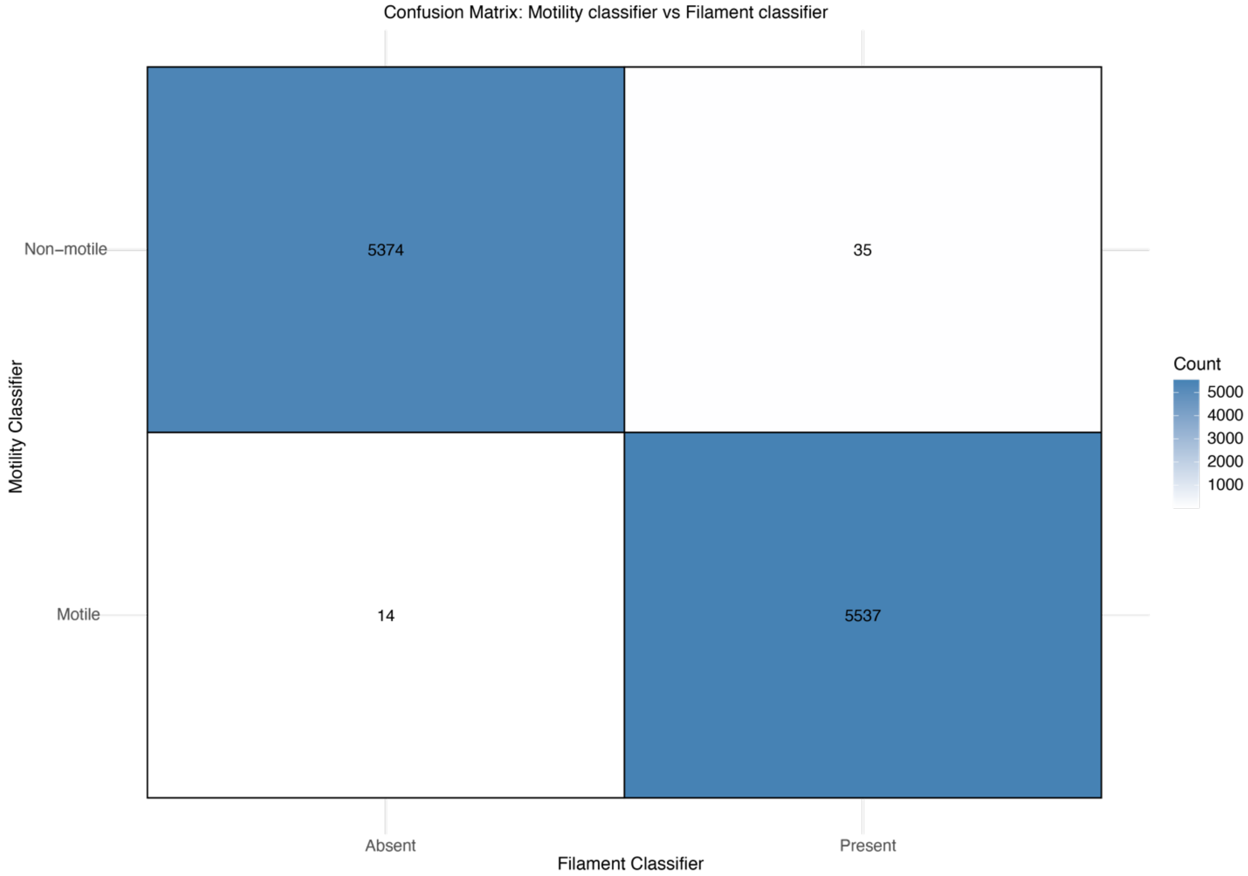

**Supplementary Fig. S11. Confusion Matrix of Filament Classifications.** This matrix quantifies presence and absence of filament proteins (FliC, FliD, FliS, FliT) for motile and non-motile clusters for 10960 species present in our dataset (full dataset with those genomes classified as partial removed). Accuracy is calculated at 99.6% as: Accuracy = (TP + TN) / (TP + TN + FP + FN)

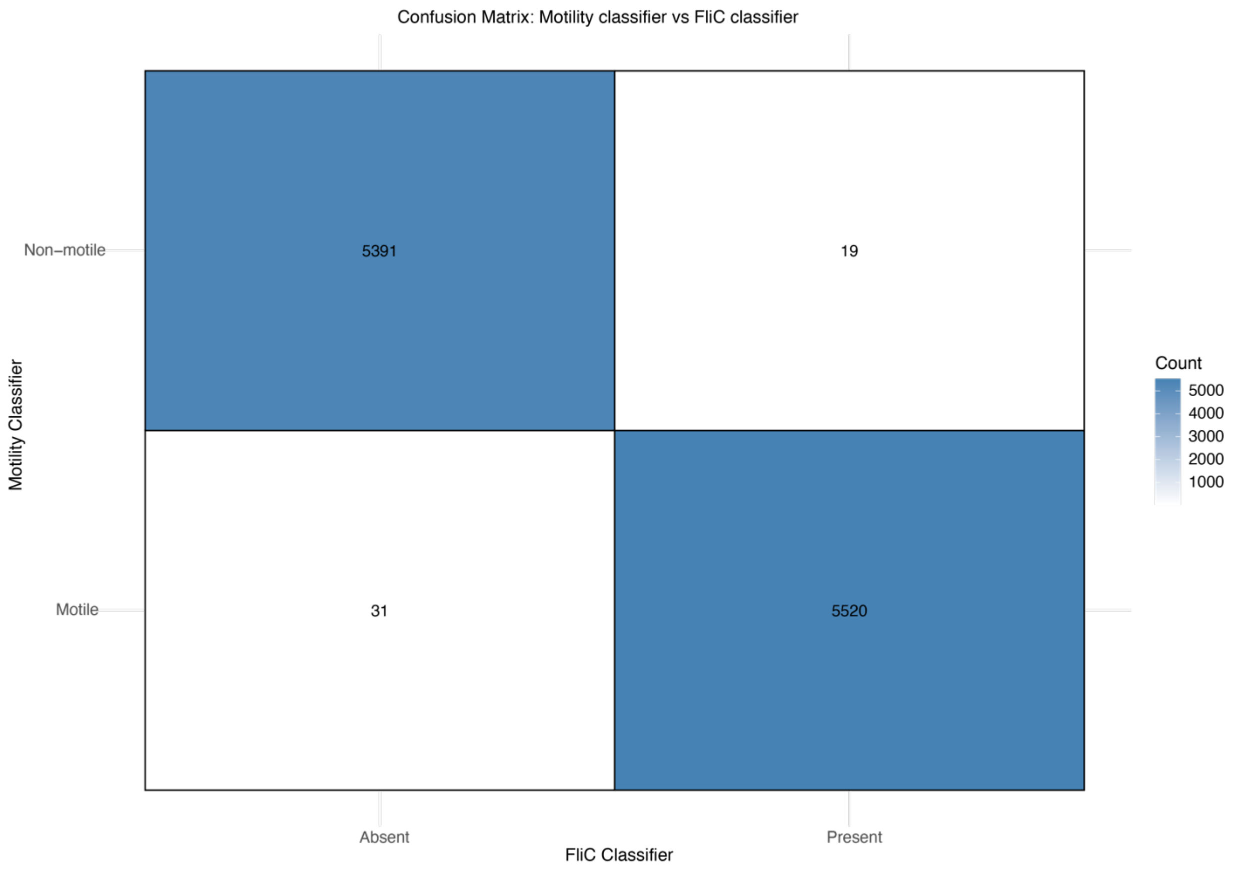

**Supplementary Fig. S12. Confusion Matrix of FliC *Classification.*** This matrix quantifies presence and absence of FliC or motile and non-motile clusters for 10960 species present in our dataset (full dataset with those genomes classified as partial removed). Accuracy is calculated at 99.5% as: Accuracy = (TP + TN) / (TP + TN + FP + FN)

| | Multistate (*Log ML*) (qMN≠qNM) | Multistate (*Log ML*) (qMN=qNM) | Log BF |
|---|---|---|---|
| Motility | -1098.5479 | -1136.74637 | 76.396876 |
| **Log BF = 2(log marginal likelihood complex model (qMN≠qNM) – log marginal likelihood simple model (qMN=qNM)** | | | |

**Supplementary Table S4:** Log Marginal Likelihood (log ML) estimates using the BayesTraits Multistate algorithms for equal and unequal rates for qMN and qNM with MCMC-and a stepping stone sampler for motility traits.

| **Protein** | | **Multistate (*LogML*) (q01≠q10)** | **Multistate (*LogML) (q01=q10)*** | **Log BF** | **Mean q10: loss** | **Mean q01:gain** |
|---|---|---|---|---|---|---|
| Filament | FliC | -1236.31922 | -1279.36122 | 86.08400 | 0.90373 | 0.25412 |
| | FliS | -1190.00789 | -1238.22466 | 96.43356 | 1.01718 | 0.27361 |
| | FliT | -801.54408 | -1032.39299 | 461.69780 | 2.89199 | 0.06330 |
| | FliD | -1201.70553 | -1249.56069 | 95.71030 | 1.02036 | 0.25515 |
| Rotor | FliG | -1501.62285 | -1507.54012 | 11.83456 | 0.90254 | 0.57065 |
| | FliM | -1282.70992 | -1311.17141 | 56.92298 | 0.86913 | 0.31444 |
| | FliN | -1326.05680 | -1365.91744 | 79.72128 | 0.97301 | 0.30650 |
| Stator | MotA | -710.72027 | -715.04653 | 8.65252 | 0.16202 | 0.45206 |
| | MotB | -1377.81916 | -1376.49928 | -2.63976 | 0.60413 | 0.60413 |
| | MotC | -133.70585 | -217.34157 | 167.27140 | 2.38561 | 0.00483 |
| | MotD | -689.31934 | -780.79494 | 182.95120 | 5.21982 | 0.27191 |
| | FliA | -716.22016 | -733.38602 | 34.33174 | 0.18116 | 0.73628 |

| | | | | | | |
|---|---|---|---|---|---|---|
| | FlhC | -658.33364 | -754.56490 | 192.46250 | 2.85142 | 0.15293 |
| | FlhD | -397.78680 | -503.70056 | 211.82760 | 2.16876 | 0.06188 |
| Type III secretion | FliH | -1583.38907 | -1589.41584 | 12.05353 | 0.78106 | 1.28892 |
| | FliI | -103.51505 | -104.33658 | 1.64305 | 0.02074 | 0.02074 |
| | FliJ | -1279.14958 | -1360.92219 | 163.54520 | 1.17715 | 0.24889 |
| | FliO | -1293.75425 | -1371.39046 | 155.27240 | 1.13792 | 0.19901 |
| | FliP | -1176.40907 | -1212.72037 | 72.62260 | 0.80795 | 0.23681 |
| | FliQ | -1195.92129 | -1232.73389 | 73.62520 | 0.83035 | 0.24063 |
| | FliR | -1185.52114 | -1224.97072 | 78.89914 | 0.83879 | 0.23415 |
| | FlhA | -1173.68779 | -1208.28648 | 69.19738 | 0.80626 | 0.24361 |
| | FlhB | -1183.56344 | -1217.71889 | 68.31088 | 0.81330 | 0.24324 |
| | FlhE | -241.15431 | -337.93076 | 193.55290 | 6.16114 | 0.04887 |
| MSP-and L rings | FliF | -1181.44873 | -1217.41895 | 71.94044 | 0.80826 | 0.23886 |
| | FlgI | -913.02227 | -1027.67297 | 229.30140 | 0.93715 | 0.05979 |
| | FlgA | -1423.41473 | -1453.98837 | 61.14728 | 0.94353 | 0.33304 |
| | FlgH | -842.18787 | -972.78843 | 261.20120 | 0.92798 | 0.04762 |
| | FliL | -1232.83343 | -1270.54440 | 75.42194 | 0.89004 | 0.26773 |
| H and T rings | FlgT | -1645.17804 | -1716.37441 | 142.39275 | 1.67847 | 0.49569 |
| | FlgO | -1136.09156 | -1181.56333 | 90.94354 | 3.12123 | 0.41400 |
| | FlgP | -669.65105 | -707.76419 | 76.22628 | 1.49630 | 0.13231 |
| | FlgQ | -60.89570 | -81.73084 | 41.67027 | 1.29738 | 0.00307 |
| | MotX | -1480.69147 | -1505.55385 | 49.72477 | 0.51793 | 1.33592 |
| | MotY | -916.49661 | -916.39461 | -0.20399 | 0.28872 | 0.28872 |
| Rod and hook | FliE | -1204.81319 | -1255.13299 | 100.63960 | 0.91200 | 0.21355 |
| | FliK | -1487.93757 | -1557.32973 | 138.78430 | 1.28489 | 0.35445 |
| | FlgB | -1808.17981 | -1849.13994 | 81.92026 | 1.54182 | 0.58505 |
| | FlgC | -1261.29225 | -1318.24910 | 113.91371 | 0.98673 | 0.22182 |
| | FlgD | -1776.49188 | -1772.77320 | -7.43737 | 0.95007 | 0.95007 |
| | FlgF | -1328.83804 | -1474.90444 | 292.13280 | 1.37494 | 0.12637 |
| | FlgG | -1409.14140 | -1538.21955 | 258.15631 | 1.56650 | 0.21522 |
| | FlgJ | -1288.13235 | -1313.68305 | 51.10140 | 0.83527 | 0.28980 |

| | | | | | | |
|---|---|---|---|---|---|---|
| | FlgE | -1306.79517 | -1367.16020 | 120.73005 | 1.02110 | 0.22166 |
| | FlgK | -1233.75053 | -1274.29683 | 81.09260 | 0.89237 | 0.26066 |
| | FlgL | -1478.32921 | -1551.66187 | 146.66532 | 1.34815 | 0.29723 |
| | FlgN | -1236.89244 | -1308.90719 | 144.02949 | 1.42868 | 0.28302 |
| Regulator | FlrA | -51.46165 | -57.63526 | 12.34721 | 0.05490 | 27.07270 |
| | FlgM | -879.56745 | -1034.52229 | 309.90969 | 1.30375 | 0.07797 |
| | FlrC | -116.83022 | -123.78185 | 13.90326 | 0.02247 | 1.13897 |
| | FliZ | -77.58300 | -141.17030 | 127.17459 | 3.55618 | 0.00827 |
| Other | FliY | -574.26291 | -629.30654 | 110.08726 | 0.10687 | 1.21131 |
| | RpoD | -70.13664 | -77.10501 | 13.93674 | 0.08113 | 28.79060 |
| | RpoN | -916.83470 | -912.49329 | -8.68281 | 0.33818 | 0.33818 |

**Supplementary Table S5:** Log Marginal Likelihood (log ML) estimates using the BayesTraits Multistate algorithms for equal and unequal rates for q01(gain) and q10(loss) with MCMC and a stepping stone sampler for individual flagellar proteins.

| **Substructure** | **Protein** | **Comparison** | **Dependent model (*Log ML*)** | **Independent Model (*Log ML*)** | **LogBF** |
|---|---|---|---|---|---|
| Filament | FliC | Motility | -1262.71280 | -2259.65168 | 1993.87776 |
| | FliS | FliC | -1609.82523 | -2428.14162 | 1636.63277 |
| | FliT | FliC | -1887.20352 | -2037.23893 | 300.07082 |
| | FliD | FliC | -1576.97481 | -2439.81757 | 1725.68551 |
| Rotor | FliG | FliC | -1921.98074 | -2737.89146 | 1631.82145 |
| | FliM | FliC | -1623.90728 | -2516.09482 | 1784.37507 |
| | FliN | FliC | -1745.42553 | -2564.63547 | 1638.41988 |
| Stator | MotA | FliC | -1619.49209 | -1947.20304 | 655.42189 |
| | MotB | FliC | -2171.08961 | -2612.84953 | 883.51984 |
| | MotC | FliC | -1341.32405 | -1369.54082 | 56.43355 |
| | MotD | FliC | -1822.04517 | -1915.64730 | 187.20426 |
| | FliA | FliC | -1913.32034 | -1950.04135 | 73.44201 |
| | FlhC | FliC | -1802.40590 | -1893.15795 | 181.50409 |
| | FlhD | FliC | -1519.01125 | -1633.49469 | 228.96686 |

| | | | | | |
|---|---|---|---|---|---|
| Type 3 secretion | FliH | FliC | -2591.42687 | -2818.24909 | 453.64443 |
| | FliI | FliC | -1330.86220 | -1339.66738 | 17.61036 |
| | FliJ | FliC | -2068.58405 | -2509.92630 | 882.68449 |
| | FliO | FliC | -1993.05548 | -2527.58780 | 1069.06463 |
| | FliP | FliC | -1469.81902 | -2414.03878 | 1888.43952 |
| | FliQ | FliC | -1505.21939 | -2429.57468 | 1848.71058 |
| | FliR | FliC | -1627.99927 | -2427.91456 | 1599.83058 |
| | FlhA | FliC | -1488.43939 | -2415.79924 | 1854.71971 |
| | FlhB | FliC | -1502.95525 | -2426.35114 | 1846.79178 |
| | FlhE | FliC | -1453.74767 | -1477.16652 | 46.83769 |
| MSPand L rings | FliF | FliC | -1490.91907 | -2418.64436 | 1855.45059 |
| | FlgI | FliC | -1579.05268 | -2149.15179 | 1140.19822 |
| | FlgA | FliC | -2375.90828 | -2658.61362 | 565.41068 |
| | FlgH | FliC | -1475.30072 | -2083.21063 | 1215.81983 |
| | FliL | FliC | -1692.94741 | -2474.08792 | 1562.28102 |
| H and T rings | FlgT | FliC | -2857.43653 | -2884.33167 | 53.79028 |
| | FlgO | FliC | -2328.72413 | -2370.49430 | 83.54033 |
| | FlgP | FliC | -1866.36753 | -1908.03772 | 83.34038 |
| | FlgQ | FliC | -1293.52206 | -1297.17991 | 7.31569 |
| | MotX | FliC | -2662.67957 | -2719.30568 | 113.25221 |
| | MotY | FliC | -2133.37752 | -2151.97865 | 37.20226 |
| Rod and hook | FliE | FliC | -1451.36643 | -2439.29563 | 1975.85840 |
| | FliK | FliC | -2082.22614 | -2712.73815 | 1261.02402 |
| | FlgB | FliC | -2207.25112 | -3044.12297 | 1673.74370 |
| | FlgC | FliC | -1549.05934 | -2504.72874 | 1911.33881 |
| | FlgD | FliC | -2254.97238 | -3013.15973 | 1516.37469 |
| | FlgF | FliC | -1984.34351 | -2566.27817 | 1163.86934 |
| | FlgG | FliC | -2121.55607 | -2647.15897 | 1051.20581 |
| | FlgJ | FliC | -2256.67354 | -2524.04003 | 534.73300 |
| | FlgE | FliC | -1597.33315 | -2539.73904 | 1884.81178 |
| | FlgK | FliC | -1455.89385 | -2470.06691 | 2028.34611 |

| | | | | | |
|---|---|---|---|---|---|
| | FlgL | FliC | -1883.79719 | -2710.82182 | 1654.04926 |
| | FlgN | FliC | -1889.40581 | -2479.26385 | 1179.71608 |
| Regulator | FlrA | FliC | -1281.63765 | -1287.77814 | 12.28098 |
| | FlgM | FliC | -1725.17754 | -2117.50901 | 784.66293 |
| | FlrC | FliC | -1347.58112 | -1353.45889 | 11.75555 |
| | FliZ | FliC | -1292.64679 | -1313.16429 | 41.03501 |
| Other | FliY | FliC | -1778.49557 | -1809.60357 | 62.21599 |
| | RpoD | FliC | -1298.88130 | -1305.03087 | 12.29915 |
| | RpoN | FliC | -2085.73875 | -2151.56623 | 131.65496 |

**Supplementary Table S6:** Log Marginal Likelihood (log ML) estimates using the BayesTraits Multistate algorithms for dependent and independent models for correlated evolution of other proteins vs FliC.
